# Surprising Threats Accelerate Evidence Accumulation for Conscious Perception

**DOI:** 10.1101/525519

**Authors:** Jessica McFadyen, Cooper Smout, Naotsugu Tsuchiya, Jason B. Mattingley, Marta I. Garrido

## Abstract

Our survival depends on how well we can rapidly detect threats in our environment. To facilitate this, the brain is faster to bring threatening or rewarding visual stimuli into conscious awareness than neutral stimuli. Unexpected events may indicate a potential threat, and yet we tend to respond slower to unexpected than expected stimuli. It is unclear if or how these effects of emotion and expectation interact with one’s conscious experience. To investigate this, we presented neutral and fearful faces with different probabilities of occurance in a breaking continuous flash suppression (bCFS) paradigm. Across two experiments, we discovered that fulfilled prior expectations hastened responses to neutral faces but had either no significant effect (Experiment 1) or the opposite effect (Experiment 2) on fearful faces. Drift diffusion modelling revealed that, while prior expectations accelerated stimulus encoding time (associated with the visual cortex), evidence was accumulated at an especially rapid rate for unexpected fearful faces (associated with activity in the right inferior frontal gyrus). Hence, these findings demonstrate a novel interaction between emotion and expectation during bCFS, driven by a unique influence of surprising fearful stimuli that expedites evidence accumulation in a fronto-occipital network.

## INTRODUCTION

The ability to predict, detect, and make decisions about danger is essential for maximising one’s chances of survival. In humans, threatening visual stimuli are detected more quickly and are more difficult to disengage from than non-threatening stimuli (Smith and Lane, 2016). Danger, however, is not always clearly visible. We must also be able to detect potential threats in visually ambiguous situations, such as when observing from a distance, in low light conditions, or when hunted by a camouflaged predator. Threats and other emotionally-salient stimuli are, indeed, more consciously accessible under difficult viewing conditions (Vieira et al., 2017). At the same time, however, our conscious perception of ambiguous visual stimuli is highly susceptible to the influence of our prior expectations, such that we tend to see what we expect to see (Hohwy et al., 2008). These prior expectations may be formed from statistical regularities in the environment that allow an organism to more efficiently respond to forthcoming stimuli. How, then, do these two neural processes interact when we are faced with a threat that we did not expect? This question has been relatively unexplored in human neuroscience, and yet it may provide important insights as to how emotion might modulate surprise signals that are propagated throughout the visual system.

Predictive coding theory suggests that our conscious perception is the result of a constant stream of hypothesis testing, where both sensory evidence and our prior expectations are integrated in a way that resembles Bayes’ rule (Rao and Ballard, 1999, Friston and Kiebel, 2009). This framework accounts for empirical evidence showing that, when sensory input is imprecise, our prior expectations or biases are weighted more heavily, consequently distorting our conscious experience (Panichello et al., 2013). For example, the perceived direction of motion is biased towards our prior expectations when motion is less coherent and thus more ambiguous (Hesselmann et al., 2010, Vetter et al., 2014). Similarly, when two different stimuli are simultaneously presented to each eye using dichoptic presentation techniques, conscious perception tends to be more stable for (Hohwy et al., 2008), and switch more rapidly to (Pinto et al., 2015), the more predictable stimulus. The expectations themselves can be established explicitly, for instance by a cue preceding the stimulus (Pinto et al., 2015, Chang et al., 2015, Meijs et al., 2018, Costello et al., 2009), or implicitly, such as by the number of stimulus presentations in the past (Barbosa et al., 2017, Aru et al., 2016, Gordon et al., 2017).

Behavioural models of perceptual decision making, like drift-diffusion modelling (DDM; Ratcliff and McKoon, 2008), have shown that prior expectations may bias the starting point of evidence accumulation such that we are predisposed towards one conclusion over another before the decision process has even begun (Barbosa et al., 2017, Mulder et al., 2012, Wiech et al., 2014, Dunovan et al., 2014, White et al., 2018, White et al., 2016). Prior expectations have also been shown to increase the drift rate of evidence accumulation (Dunovan et al., 2014, White et al., 2016) and may lower the threshold for awareness (De Loof et al., 2016). Other components of the decision making process, such as sensory processing and/or motor response execution (known collectively as non-decision time; Ratcliff and McKoon, 2008) have also been shown to speed up with prior expectations (Jepma et al., 2012) or when stimuli are self-relevant (Macrae et al., 2017).

Like predictable events, threatening stimuli are also prioritised for conscious access (Otten et al., 2017). For example, fearful faces, snakes and spiders (Gomes et al., 2017), and fear-conditioned stimuli (Gayet et al., 2016) are consciously perceived earlier than neutral stimuli during breaking continuous flash suppression (bCFS; Tsuchiya and Koch, 2005, Jiang et al., 2007). Fearful stimuli have been shown to increase the rate of evidence accumulation (Tipples, 2015) even when unconsciously-presented (Lufityanto et al., 2016), as well as bias the starting point towards threat (Zaman et al., 2017). There has, however, been very little investigation into how the prioritisation of fearful stimuli for conscious access might be influenced by prior expectations.

We propose three testable hypotheses for how prior expectations might influence conscious access to suppressed threatening and neutral stimuli. The first is the **Emotional Exaggeration Hypothesis**, which proposes that. the effect of expectation on conscious perception is exaggerated for emotional stimuli. Previous studies have found that surprise-related evoked potentials are larger and earlier for emotional than neutral stimuli (Vogel et al., 2015, Kovarski et al., 2017, Chen et al., 2017). If the effect of expectation is even *larger* for emotional stimuli, as this suggests, then we might expect that earlier conscious perception of expected than unexpected stimuli (after an initial period of unawareness) is even more extreme for emotional stimuli. Indeed, previous inattentional blindness research suggests that both emotional and neutral stimuli are missed equally as often when they are unexpected but emotional stimuli are detected more frequently than neutral stimuli when expected (Beanland et al., 2017, Wiemer et al., 2013).

As an alternative to the Exaggeration Hypothesis (where the effect of threat on expectation is synergistic), we might consider a **Survival Hypothesis**, such that threat negates or reverses the effect of expectation on conscious perception. This captures the notion that, even in situations where a threat is unexpected, it is still vital (or, arguably, even *more* vital) that we can rapidly respond (Den Ouden et al., 2012). Studies on attentional capture and inattentional blindness have found that, while neutral stimuli evoke slower responses when they are unexpected, threatening stimuli elicit the fastest responses in visual search tasks regardless of prior expectations (Aue et al., 2016, Aue et al., 2013). Other research, however, has shown that novelty detection and attentional biases towards threat are *enhanced* in contexts where threats are unpredictable (Garcia-Garcia et al., 2010, Aue and Okon-Singer, 2015, Bar-Haim et al., 2007, Notebaert et al., 2010). Additionally, unexpected threats are more frequently detected than unexpected neutral images (New and German, 2015) and evoke stronger physiological responses (Wiemer et al., 2013), even under high perceptual load (Gao and Jia, 2017). It is thought that subcortical ‘survival circuits’ involving the amygdala facilitate rapid modulation of conscious perception, such as faster threat detection in visual search and attentional blink paradigms (Tamietto and De Gelder, 2010, Mitchell and Greening, 2012). Hence, such facilitation may circumvent or interact with the influence of top-down expectations, resulting in earlier conscious access to emotional stimuli that is unmodulated or hastened by surprise (Hohwy, 2012).

In contrast to the first two hypotheses that predict a synergistic (**Emotional Exaggeration Hypothesis**) or antagonistic (**Survival Hypothesis**) interaction between threat and expectation, a third possibility is that threat and expectation do not interact. Some inattentional blindness research has found no advantage of unexpected threats versus non-threats in entering awareness (Calvillo and Hawkins, 2016, Beanland et al., 2017). Hence, we also considered the **Additive Hypothesis**, which is that both expectation and emotional content independently accelerate conscious perception without interacting.

To test the three hypotheses above, we conducted two bCFS experiments and one control experiment (see **Supplementary Materials**). In Experiment 1, we established how expectation interacts with threat in bringing stimuli into conscious perception. In Experiment 2, we adapted the design of Experiment 1 to incorporate EEG so that we could observe the spatio-temporal maps of neural activity underlying the effects of emotion and expectation during bCFS. We also conducted drift diffusion modelling (DDM) to examine which parameters of the decision-making process explained the differences in response time between conditions and how this was reflected in neural activity. DDM has been used in previous studies investigating consciousness that equate the upper decision boundary to the threshold for awareness (De Loof et al., 2016, Kang et al., 2017). Here, response times in the bCFS paradigm reflected a perceptual decision (whether the face was rotated clockwise or anticlockwise) that required conscious perception (Kang et al., 2017). We investigated how the rate of evidence accumulation (drift rate; *v*), sensory processing and/or motor execution (non-decision time; *t*_*0*_), and the decision boundary (*a*) might be influenced by threat and expectation.

## METHODS

### Participants

We recruited 30 participants for Experiment 1 and 33 participants for Experiment 2 through the University of Queensland’s Participation Scheme, which draws from adults within the local community. Our sample for Experiment 1 consisted of 13 males and 17 females aged between 18 and 27 years (*M* = 21.50, *SD* = 2.36). For Experiment 2, we removed 2 participants for having insufficient trial numbers (see **Analysis** section), leaving 17 males and 14 females aged between 18 and 28 years (*M* = 22.00, *SD* = 2.08). All participants reported having normal short- and long-distance vision without the need for glasses or contact lenses. Participants were compensated AUD$20 per hour for their time and provided written consent. This study was approved by the University of Queensland’s Human Research Ethics Committee.

### Stimuli

We collected face stimuli from a variety of experimentally-validated databases to maximise the number of unique stimuli presented throughout the experiment. This included 24 images from the Amsterdam Dynamic Facial Expressions Set (ADFES; van der Schalk, Hawk, Fischer, & Doosje, 2011), 132 images from the Karolinska Directed Emotional Faces set (KDEF; Lundqvist, Flykt, & Öhman, 1998), 52 images from the NimStim set (Tottenham et al., 2009), and 58 images from the Warsaw Set of Emotional Facial Expression Pictures (WSEFEP; Olszanowski et al., 2015). The final selection consisted of 267 images of Caucasian adults (66 females and 71 males) displaying either a neutral or fearful facial expression. We cropped the hair, neck, and shoulders from all face stimuli (see **Fig. 1**). We then centred the faces within a 365 × 365 pixel square with a grey background for Experiment 1 and a black background for Experiment 2 (to maximise the visually-evoked EEG response to the face, due to the greater contrast difference for greyscale stimuli fading in from black than grey). We equated luminance and root-mean square contrast (of pixels in the entire image, including the face and grey background) across all images using the SHINE toolbox (Willenbockel et al., 2010), such that they did not differ significantly between neutral and fearful faces (luminance: neutral = 125.080, fearful = 124.681, *t*(130) = 1.954, *p* = .106; contrast: neutral = 125.903, fearful = 125.472, *t*(130) = 2.038, *p* = .088; Bonferroni-corrected for two comparisons).

**Figure 1.**
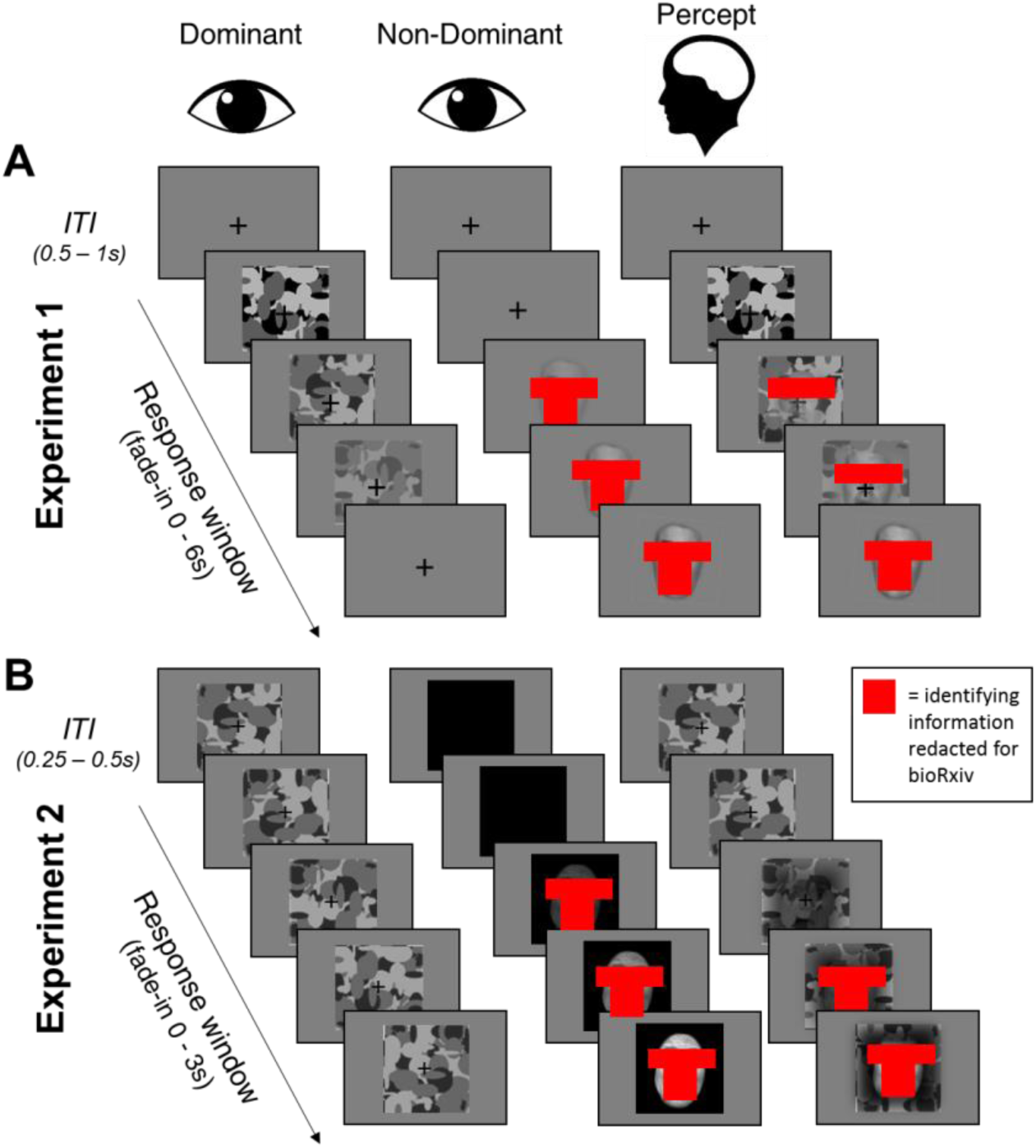
Schematic for the basic paradigm across experiments. In Experiment 1 (**A**), the face linearly faded from 0% to maximum contrast over 6 s, while the mask did the opposite in the non-dominant eye. Experiment 2 (**B**) was the same except that the fade period was 3 s, the mask remained at maximum contrast, and the face background was black rather than grey. In both experiments, participants perceived the mask at first, followed by a period of mixed percept of the mask and the face (as shown in the ‘Percept’ column). For Experiment 1, the trial ended upon response, whereas for Experiment 2, the trial always ended at 3 s regardless of response. The inter-trial interval (ITI) was jittered between 0.5s and 1s for Experiment 1, and between 0.25 and 0.5s for Experiment 2 (the mask was also shown throughout the ITI for Experiment 2). Note that identifying information (facial features hidden by red boxes) has been redacted from this preprint.

We used Mondrian images to mask the stimuli (see **Fig. 1**). These images were made using code available online (http://martin-hebart.de/webpages/code/stimuli.html; as used in Stein, Seymour, Hebart, & Sterzer, 2014). The Mondrian images were presented at 125% of the size of the face stimuli to ensure that faces were sufficiently well-masked.

### Procedure

#### Dichoptic presentation set-up

After completing the consent form, participants completed the self-report 40-item State-Trait Anxiety Inventory (STAI; Spielberger, Gorsuch, & Lushene, 1970). We then determined the participants’ ocular dominance using the Miles Test (Miles, 1930). Participants then sat approximately 1.1m (Experiment 1) or 0.55m (Experiment 2) from a 22” LCD monitor (1980 × 1020 resolution) with a black screen divider placed in front (see **Supplementary Fig. 1**). For Experiment 1, the participant positioned their head in a chin and head rest, to which prism lenses (12 prism diopters, base out) were attached and secured with a foam strap. For Experiment 2, stereoscopic mirrors were used instead of prism lenses as they were faster to set up. Both methods result in dichoptic presentation (see **Supplementary Fig. 1**).

In both experiments, participants completed a short calibration task (using placeholder stimuli the size of the mask) and the apparatus was adjusted (i.e. angle of mirrors/prism lenses, computer monitor height, etc.) to ensure that the stimuli presented to each eye were perceived to be in the same location in space (i.e. completely overlapping in the centre of field of vision) and that only *one* stimulus could be perceived with each eye. Note that in Experiment 1, an eye tracker was also used to ensure that participants did not close one eye during the experiment (which would interrupt the interocular suppression).

#### Behavioural paradigm

Each trial began with the mask presented at 100% contrast to the participant’s dominant eye and a face stimulus presented at 0% to the other eye (see **Fig. 1**). In Experiment 1, the stimuli were set to fade over a period of 6 s, with the mask fading out from 100% to 0% contrast and the face fading in from 0% to 100% contrast. Experiment 2 was the same, except the time period was reduced to 3 s (to reduce experiment length and increase the number of trials) and the mask contrast was fixed at 100% (to avoid an onset effect in the EEG signal). In both experiments, the face images were pseudo-randomly rotated 5° clockwise or counter-clockwise. Participants were instructed to click the left or right mouse button as soon as they could perceive the orientation of the face. Participants were told to maintain accuracy as close to 100% as possible while also making the responses as fast as possible. In Experiment 1, trials ended upon response (if responses were over 6 s, the face remained at 100% and the mask at 0% until response), whereas in Experiment 2, trials always ended after 3 s regardless of response. Between trials, a fixation cross was presented at the centre of each left and right image frame. The duration of the inter-trial interval (ITI) jittered randomly between 0.5 and 1 s at a step of 0.1 s for Experiment 1, and between 0.25 to 0.50 s at a step of 0.05 s for Experiment 2.

There were eight blocks in Experiment 1 and fourteen blocks in Experiment 2. In both experiments, participants were informed that some blocks would contain more of one emotional expression than others but that this was irrelevant to their task (i.e. they were to respond to every face they saw, regardless of emotion). Half of the blocks contained predominantly (83%) neutral faces while the other half of the blocks contained predominantly fearful faces. The dominant emotional expression was indicated at the beginning of each block by a 5 s presentation of the word “Neutral” or “Fearful”. Neutral and fearful blocks were alternated, with the starting block emotion counterbalanced across participants. There were 90 trials per block and each block began with at least two trials for the predominant emotion. The presentations of rare and unexpected (17%) emotional faces were thereafter spaced apart by 2 to 7 trials (following a Gaussian distribution). There were 720 total trials for Experiment 1 (300 expected and 60 unexpected trials per neutral/fearful expression) and 1,260 total trials for Experiment 2 (525 expected and 105 unexpected trials per neutral/fearful expression).

#### Behavioral titration procedure for EEG recording (Experiment 2)

In Experiment 2, participants completed a titration task while the EEG cap was set up. The purpose of the titration task was to ensure that responses could be made on the majority of trials (e.g. participants more susceptible to masking effects might take longer than the 3-second trial window to consciously perceive the face, thus making less responses overall). The titration task consisted of four blocks: two neutral-dominant and two fearful-dominant blocks in an alternate order, with the starting block counterbalanced across participants. Each block contained 90 trials, with 83% dominant emotion presentations and 17% rare emotion presentations. All aspects of the titration trials (e.g., stimuli, timing) were the same as the trials in Experiment 2 (see **Fig. 1**).

The titration began with the mask at low contrast relative to the face (100% face, 0% mask). Using the Palamedes toolbox (Prins and Kingdom, 2009), contrast was adjusted per trial, such that if the response was faster than 2 s, the next trial’s face contrast would be decreased and mask contrast increased (hence, mask contrast always equalled 1 minus the face contrast), and vice versa for responses slower than 2 seconds. Thus, the face and mask contrasts were adjusted so that conscious breakthrough occurred approximately two thirds of the way into each trial for each participant, accommodating for individual differences in sensitivity to interocular suppression. The stepwise function used for these trial-by-trial adjustments began with 10% contrast adjustments, which were reduced by 2% each time a reversal (i.e. a change in response type; fast to slow, or slow to fast) was made. After 4 reversals, contrast adjustments were fixed at 2%. These staircases were constructed independently for the first two blocks of titration trials (one neutral, one fearful), resulting in one ending set of contrasts per emotion. This value was used as the starting point for the *second* block of each dominant emotion, giving a fine-tuned contrast set built across two blocks of 90 trials each per neutral and fearful block type. The neutral-dominant and fearful-dominant contrast sets were then averaged together, giving face contrast values ranging from 53.23% to 91.68% (*M* = 76.75%, *SD* = 10.25%) across participants (mask contrast values were equal to 1 minus the face contrast). Each participant’s final titrated face contrast value was used for all face stimuli (neutral or fearful, in any block type) presented in the main experiment, where faces faded in from 0% to the titrated value over 3 seconds. Although the first two participants did not complete the titration task, their mean reaction times throughout the experiment were 1.803 s and 2.288s, respectively, and so they were included in further behavioural and EEG analyses.

### EEG recording

#### EEG acquisition

Neural activity was continuously recorded using a BioSemi Active Two 64 Ag-AgCl electrode system (BioSemi, Amsterdam, Netherlands) throughout the 14 experiment blocks. Participants were fitted with a nylon cap containing 64 Ag/AgCl scalp electrodes positioned according to the international 10-20 system. Continuous data were recorded using BioSemi ActiView software (BioSemi, 2007), referenced to the standard BioSemi reference electrodes, filtered online (0.01 to 208 Hz amplifier band pass filter), and then were digitised and stored at a sampling rate of 1024 Hz with 24-bit A/D conversion. We measured horizontal and vertical electrooculograph (EOG) signals with flat biploar Ag/AgCl electrodes. The experiment was conducted in an electrically-shielded Faraday cage to minimise noise and all data was recorded with electrode impedance levels under 25 µV.

#### EEG preprocessing

All preprocessing was done via MATLAB 2016a (MathWorks). Data were imported into SPM12 (Wellcome Trust Centre for Neuroimaging, London). The data were then re-referenced to the average across all 64 EEG scalp channels and the pairs of vertical and horizontal EOG electrodes were referenced to each other. Noisy channels were interpolated using FieldTrip (Oostenveld et al., 2011). Eyeblinks were marked using the vertical EOG and the associated spatial confounds were corrected using SPM12’s signal-space projection (SSP) method. The data were then bandpass filtered between 0.1 and 40 Hz and epoched into −0.1 to 3 s segments around stimulus onset for event-related potential (ERP) analyses. Each epoch was baseline-corrected (mean amplitude subtraction) using the −0.1 to 0 s period pre-stimulus-onset. Trials with incorrect responses or response times more than 3 standard deviations from the mean (within-participant, collapsed across each condition) were excluded, so that the EEG data represented the typical responses for each participant. The data were then robust-averaged (i.e. the contribution of each trial to the average, iteratively weighted by noise level; Wager et al., 2005) and the bandpass filter and baseline correction were re-applied. Finally, in order to conduct statistical parametric mapping (Penny et al., 2011) in SPM 12, we converted the robust-averaged ERPs into three-dimensional images (*x* and *y* space, *ms* time) and smoothed them with a 12 mm x 12 mm x 12 ms FWHM Gaussian kernel to accommodate for intersubject variability.

### Analysis

#### Behavioral preprocessing

Within each participant’s data, we first removed responses faster than 500 ms (e.g. accidentally pressing the mouse button too quickly; median = 0, range = 0 to 90 trials removed per participant for Experiment 1 and median = 4, range = 0 to 87 trials for Experiment 2). Accuracy on the orientation task was near ceiling in both experiments (mean and standard deviation of accuracy for Experiment 1: expected neutral = 98.0% ± 1.8%, unexpected neutral = 97.5% ± 2.2%, expected fearful = 96.9% ± 2.6%, unexpected fearful = 97.9% ± 2.5%; Experiment 2: expected neutral = 94.5% ± 5.9%, unexpected neutral = 95.1% ± 4.0%, expected fearful = 94.2% ± 5.3%, unexpected fearful = 94.9% ± 4.9%). We only entered the correct trials into our response time analysis and EEG analysis. We removed responses more than five standard deviations from the mean (e.g. lapse in attention to experiment; approximately *M* = 1, *SD* = 1 trials removed per participant for Experiment 1 and no outliers detected in Experiment 2) collapsed across conditions. For Experiment 1, the average trial counts were 291 for expected neutral (225 to 300), 292 for expected fearful (277 to 300), 58 for unexpected neutral (54 to 60), and 58 for unexpected fearful (45 to 60) faces. For Experiment 2, two participants had less than 80 trials in at least one condition (due to very slow responses that went into the next trial) and thus were deemed to have insufficient data for EEG analysis. After removing these two participants, the average trial counts for 31 participants were 493 for expected neutral (379 to 522), 496 for expected fearful (429 to 523), 99 for unexpected neutral (80 to 104), and 99 for unexpected fearful (84 to 105) faces.

#### Linear mixed effects modelling

To investigate differences in response time between conditions, we entered the data into a hierarchical series of linear mixed effects (LME) models using the “lme4” package (Bates et al., 2014) in R v3.4.3 (Team, 2014). The LME is an extension of linear regression that estimates both fixed and random effects (Gelman and Hill, 2006). This approach allowed us to encapsulate all data from all participants (rather than taking a mean or median response time per participant) and to account for different trial numbers per experimental condition (Baayen et al., 2008). LME also allowed us to model fixed and random effects of no interest to maximise statistical power (Barr et al., 2013). These included the fixed effects of participant gender and block order (i.e. whether participants were assigned a neutral or fearful block first), and the random effects of participant (as the experiment was a repeated-measures design), trial number (indicating fatigue and/or learning effects across the duration of the experiment) and anxiety (summed state/trait score from the STAI questionnaire). Summed anxiety scores (which can range from 40 to 160) for Experiment 1 ranged from 46 to 107 (*M* = 69.97, *SD* = 15.90) and for Experiment 2 ranged from 40 to 108 (*M* = 71.94, *SD* = 16.02). We constructed a series of 5 hierarchical models, with the first encapsulating just the effects of no interest and then each subsequent model incorporating an additional effect of interest (in this case, the main effects of emotion, expectation, and their interaction). To test for significant differences between these models, we performed likelihood ratio tests.

#### Drift diffusion modelling

To better elucidate the mechanism by which expectation influences response times to neutral and fearful faces, we employed drift diffusion modelling (Ratcliff, 1978). DDM depicts binary decision-making as a stochastic process, whereby evidence gradually accumulates (with added noise) from a starting point (*z*) towards one of two thresholds (*a*). Several other parameters influence the resultant response, such as the drift rate (*v*; the rate of evidence accumulation) and non-decision time (*t*_*0*_; the duration of stimulus encoding; Ratcliff and McKoon, 2008). We focused on the parameters for the decision threshold (*a*), drift rate of evidence accumulation (*v*), and the non-decision time for stimulus encoding (*t*_*0*_). Specifically, we were interested in how these parameters might differ between emotion and expectation conditions.

For the parameter optimisation, rather than allowing all three parameters of interest to vary per condition (resulting in 12 parameters instead of 3), we constructed a series of eight models to see which combination of condition-specific parameters (e.g. 4 *a* parameters, one per condition, with 1 *v* and 1 *t*_*0*_ parameter for all the data, versus 4 *v* and 4 *t*_*0*_ parameters, one per condition, with 1 *a* parameter for all the data, etc.) best explained the group data (all trials pooled across participants). The eight models were: 1) no condition-specific parameter optimisation, 2) *a*, 3) *v*, 4) *t*_*0*_, 5) *a* and *v*, 6) *a* and *t*_*0*_, 7) *v* and *t*_*0*_, and 8) *a, v*, and *t*_*0*_. We pooled the data across all participants so that we maximised our power, due to the relatively lower number of incorrect trials (correct responses: count = 36,802, count per condition = 3,055 to 15,390, mean RT = 1.844 ± 0.423; incorrect responses: count = 801, count per condition = 52 to 356, mean RT = 1.887 ± 0.501).

We used the *fast-dm* software (Voss and Voss, 2007) to optimise the parameters using maximum likelihood estimation. For the estimation, we fixed four variables based on the design features of our task. First, we fixed *z* to 0.5 because face orientation was randomised and thus participants could not be biased towards a correct or incorrect orientation before the trial had begun. Second, we fixed differences in speed of response execution to 0 because we expected motor responses to be relatively uninfluenced by expectation or emotion. Third, we fixed inter-trial variability of *z* and *v* to 0 because there were low trial numbers for the incorrect responses to reliably estimate inter-trial variability. This left *a, v, t*_*0*_, inter-trial variability in *t*_*0*_(as recommended by Voss and Voss, 2007), and percentage of contaminants (i.e. guesses) as free parameters that could vary either generally or condition-specifically (expected/unexpected and neutral/fearful faces), depending on the model. We then compared the minimised log likelihood across all eight models, as well as the AIC (criteria for best model ≥ 3; Raftery, 1995) to account for models with more parameters, to see which parameter optimisation set-up resulted in the best explanation of the data. We also used the Kolmogorov-Smirnov test statistic (*p*) as a measure of model fit, where *p* > .05 indicates sufficient goodness-of-fit. We adjusted *p* for models with condition-specific free parameters by calculating *p*^1-^_*k*_, where *k* is the number of conditions (Voss and Voss, 2007).

After establishing the best parameter optimisation set-up across all participants, we took the winning model architecture and conducted parameter optimisation on each participant so that we could statistically compare differences in condition-specific parameters using a 2 × 2 repeated-measures ANOVA (between emotion and expectation).

### EEG analysis

#### Robust-averaged ERPs

We conducted general linear model (GLM) analyses in SPM 12 on the robust-averaged ERPs per condition using each participant’s smoothed 3D images. Each GLM consisted of a 2 (expected, unexpected) × 2 (neutral, fearful) repeated-measures design, where we investigated the main effects of emotion and expectation, and their interaction. Each participant’s gender, block order (i.e. whether the experiment began with a predominantly neutral or fearful block), and anxiety (summed state and trait scores) were also entered as covariates of no interest and mean-centred. We compared four variations on this GLM, each of which incoporated the behavioral data to explain the observed ERP data in a distinct way. The first GLM consisted of the above design without any additional components. The second GLM included participant’s average response time per condition as a covariate of interest. The third and fourth GLMs were ‘model-based’ GLMs (O’doherty et al., 2007), such that they included covariates for either the *v* or *t*_*0*_ parameter estimates per condition derived from DDM per condition. For each of these GLMs, the resultant SPMs were corrected for multiple comparisons according to Random Field Theory (Worsley, 2006). Given the large number of multiple comparisons in this particular experimental design (due to the long epoch duration of 3 seconds – 3,072 samples), we applied a small volume correction so that we only examined results across the scalp from 0 to 2 seconds as we were primarily interested in neural activity *preceding* response time (average response times = 1.844 s, *SD* = 0.423 s). Only clusters with *p* <.05 family-wise-error (FWE) corrected were considered.

#### Source reconstruction

For cortical source reconstruction from our EEG data, we used the Multiple Sparse Priors (MSP) method (Friston et al., 2008) implemented in SPM12. This method is a Bayesian solution to the EEG inverse problem that puts certain constraints (i.e. priors) on likely sources of observed EEG activity, including that the sources are likely multiple and sparse. First, a head model was constructed for each participant’s EEG data using a canonical T1 image provided by SPM12 and estimated using a single-sphere Boundary Element Model (Mattout et al., 2007). We then optimised the inversion process by simultaneously inverting the data for each condition across all 31 participants (i.e. group inversion), thus assuming that the responses in all participants (who each completed the same experimental task) should be explained by the same set of sources (Litvak and Friston, 2008). After group inversion, we then extracted cortically-smoothed images over space (8 × 8 × 8 mm FWHM Gaussian kernel) of the estimated source activity per condition across several time windows of interest. The first window was 0 to 2 seconds (similar to the ERP analysis described above). The second, third, and fourth were 500 ms bins from 0.5 to 1 s, 1 to 1.5 s, and 1.5 to 2 s, allowing us to observe how source activity changed over time.

Our first statistical analysis was a 2 (emotion) × 2 (expectation) full factorial GLM, where the levels of each factor were dependent and the variance was assumed to be unequal. We entered in gender, block order, and anxiety as mean-centred covariates of no interest and then examined the main effects and interactions (*F* tests). If significant, these were followed up with *t*-tests to examine the specific direction of each effect. We then conducted separate GLMs to examine how cortical sources evolved over time as a function of two key DDM parameters: drift rate (*v)* and non-decision time (*t*_*0*_), which were found to vary in the winning model from the DDM group optimisation stage described earlier. Hence, there were six separate GLMs (2 parameters (*v* and *t*_*0*_) × 3 time windows – 500 ms each). The design of each was the same as the first (i.e. 2 × 2 full factorial with 3 covariates of no interest) but had an additional DDM covariate of interest (either *v* or *t*_*0*_ value per condition, per participant). To follow up the behavioural results we observed earlier, we computed specific *t* contrasts. Specifically, for drift rate (*v*), we found behavioural evidence that *v* is interactively influenced by emotion and expectation, and so we tested if the *v* parameter covaried with interactive effects at the source level (that is, greater activity for unexpected than expected fearful faces, compared with neutral faces). Similarly, we found behavioural evidence that non-decision time (*t*_*0*_) was influenced by expectation, and so we tested if the *t*_*0*_ parameter covaried with stronger (contrast 1) or weaker (contrast 2) source activity for surprise. For all GLMs, the SPM Anatomy Toolbox was used to identify the significant sources (Eickhoff et al., 2005), *p* < .05 cluster-level family-wise-error-corrected.

## RESULTS

### Prior expectations speed up breakthrough of neutral but not fearful faces

Experiment 1 was our first investigation into how emotion and expectation might interactively influence breakthrough times in a bCFS paradigm. Using LME to model all trial data (accounting for inter-participant variance, gender, block order, trial number, and anxiety score; see **Fig. 2C**), we discovered that the interaction model (response time ∼ emotion × expectation + gender + block order + random effect of subject + random effect of anxiety + random effect of trial number) was the highest performing (χ^2^ = 9.282, *p* = .002; see **Fig. 2A**) compared to the other nested models (i.e. the null model, emotion model, expectation model, and emotion + expectation model). The least-squares means (in seconds) predicted by the winning model revealed a significantly slower estimated response time for unexpected (*M* = 3.355; *95% CI* [2.782 3.927]) than expected (*M* = 3.443; *95% CI* [2.889, 4.047]) neutral faces (*p* = .0001), while there was no significant difference between expected (*M* = 3.158; *95% CI* [2.610, 3.754]) and unexpected (*M* = 3.148; *95% CI* [2.601, 3.759]) fearful faces (*p* =.954; see **Fig. 2B**), which were faster than neutral faces overall.

**Figure 2.**
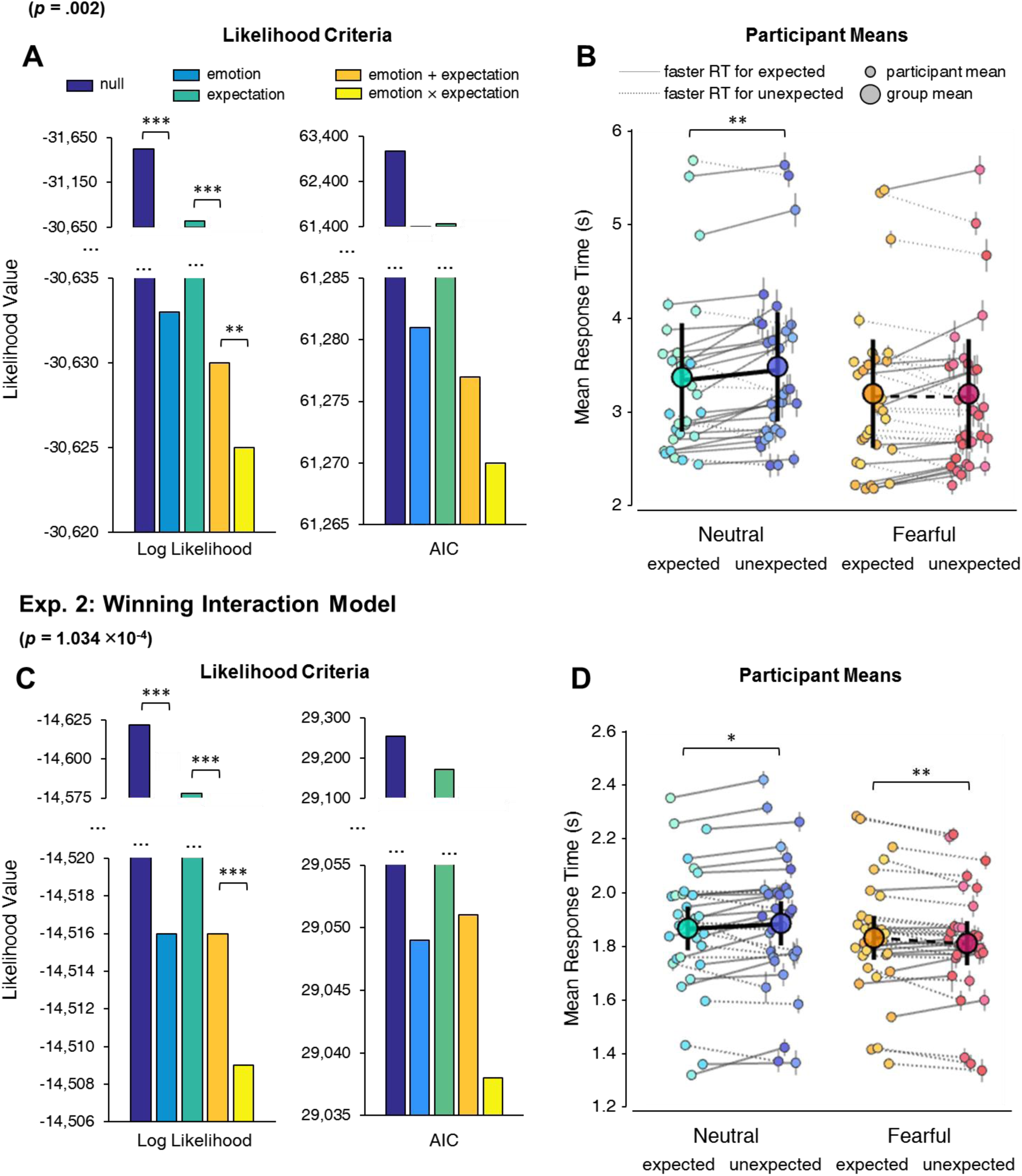
Winning interaction models from Experiments 1 and 2. The LME results are displayed for Experiment 1 (**A** and **B**) and Experiment 2 (**C** and **D**). **A** and **C** display the likelihood of each model as given by the log likelihood and the Akaike information criteria (AIC) during likelihood ratio estimation (both measures are better when the height of the bars are lower). Asterisks (* *p* < .05, ** *p* < .01, *** *p* < .001) indicate the significance of log likelihood ratio tests between models, and arrows point towards the smallest AIC values. **B** and **D** display each participant’s mean response time (*y*-axis) per condition (*x*-axis), as represented by the smaller dots (error bars represent standard error across trials). The lines connect expected and unexpected conditions for a single participant, with solid lines indicating faster responses to expected faces and dashed lines indicating faster responses to unexpected faces. The least-squares mean estimated across the entire dataset by the winning model is represented by the larger circles, where error bars represent 95% confidence interval. Asterisks indicate the significance of the simple effects of prediction for each emotion (as given by least-squares means).

We conducted Experiment 2 on 31 new participants to investigate whether patterns of neural activity unfolded differently over time between emotion and expectation conditions during interocular suppression. We modified the paradigm from Experiment 1 to accommodate the EEG recording (see **Methods** for details). We replicated the main behavioural results from Experiment 1 using this modified version of the task on a new group of participants. The interaction model was, again, the highest performing model relative to all others (χ^2^ = 15.075, *p* = 1.034 × 10^−4^; see **Fig. 2D)**. The least-squares means (in seconds) predicted by the winning model revealed a significantly slower estimated response time for unexpected (*M* = 1.863; 95% CI [1.782, 1.944]) than expected (*M* = 1.882; 95% CI [1.800, 1.964]) neutral faces (*p* = .010). This time, however, the least-square means were significantly faster for unexpected (*M* = 1.808; 95% CI [1.726, 1.890]) than expected (*M* = 1.828; 95% CI [1.747, 1.909]) fearful faces (*p* = .007; see **Fig. 2E**).

Overall, the behavioural results from Experiments 1 and 2 demonstrate that implicitly-learned expectations for emotional expression accelerated reponses to neutral faces, while there was either no effect (Experiment 1) or the *opposite* effect (Experiment 2) on fearful faces. Hence, these results favour the Survival Hypothesis, such that both expected and unexpected fearful stimuli were prioritised for conscious access. This is in contrast with the Additive Hypothesis (that there would be an equal influence of prior expectations on neutral and fearful faces) and the Emotional Exaggeration Hypothesis (that there would be an even larger effect of expectation on fearful than neutral faces).

### Prior expectations shorten non-decision time and unexpected threat accelerates evidence accumulation

We used drift diffusion modelling to explain the response time patterns (i.e. faster for expected than unexpected neutral faces, while the opposite was true for fearful faces). We conducted this analysis on the behavioural data from Experiment 2, rather than Experiment 1, because Experiment 2 contained considerably more trials and also allowed us to relatethe resultant decision parameters to the EEG data. In an initial group-level parameter optimisation step (see **Methods** for details), we established that the data overall were best explained when the drift rate (*v*) and non-decision time (*t*_*0*_) were free to vary per condition (giving four condition-dependent values for each parameter), while the threshold (*a*) parameter was free to vary across all the data generally – this was model 7 (*v, t*_*0*_; see **Fig. 3A**). This model’s AIC was sufficiently lower (23.7) than the next best model, model 3 (*v*), and sufficiently fit the observed data (Kolmogorov-Smirnov test statistic: null model *p* = 0.111 and winning model *p* = 0.593, where an adequate model fit is indicated by *p* > .05).

**Figure 3.**
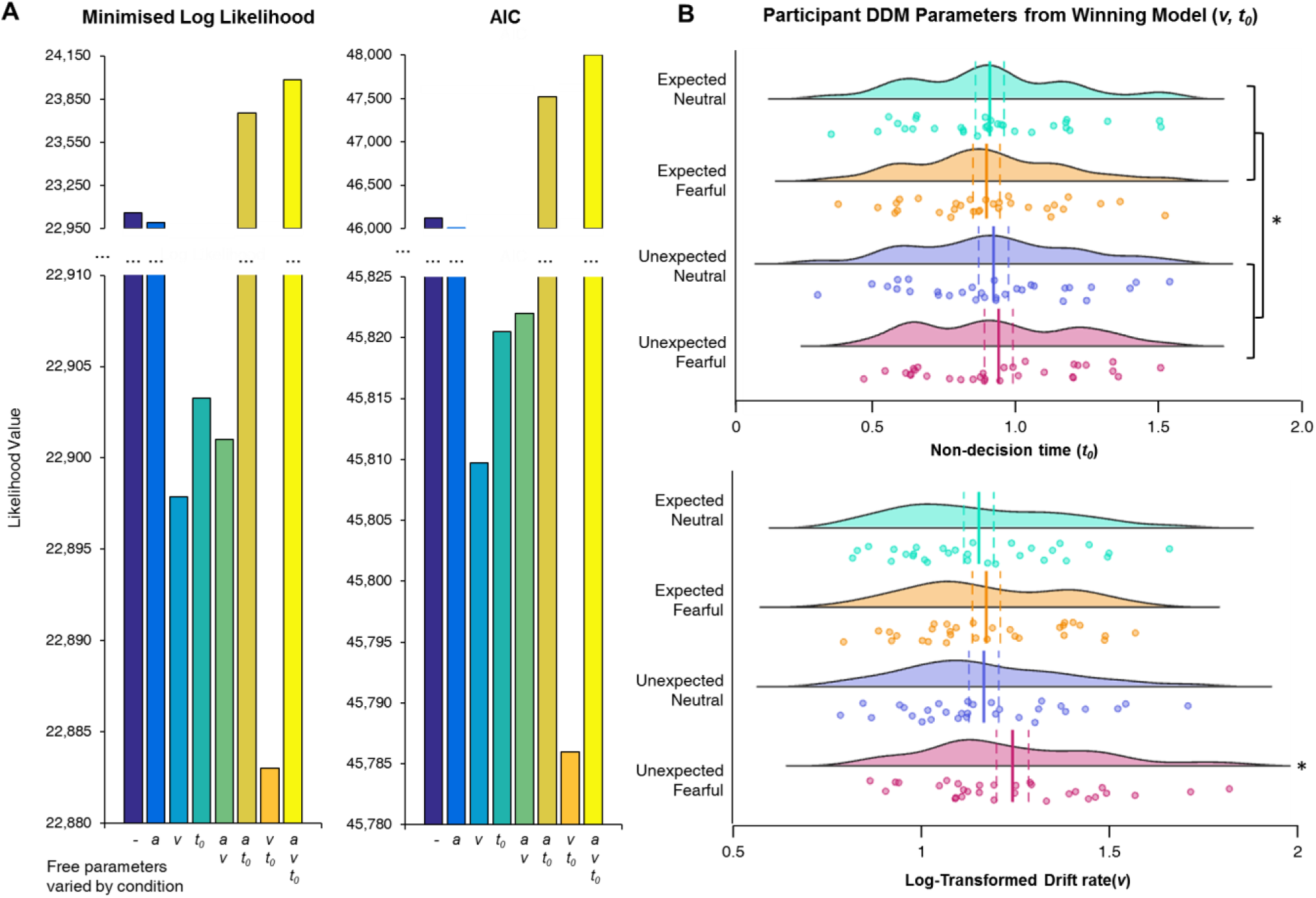
Drift diffusion modelling group-level model comparison and participant-level parameter comparisons. **A)** The results of the group-level parameter optimisation are shown. The minimised log likelihood (left) and AIC (right) values are shown for each of the eight models, where the parameters that were free to vary per condition are indicated along the *x* axis. Arrows indicate the model with the lowest value. **B)** The estimated parameter values for *v* (top) and *t*_*0*_ (bottom) are shown per participant. These ‘raincloud’ plots illustrate the distribution of data (histogram) and the individual data points (each participant). The mean is indicated by vertical solid lines and standard error is represented by dashed vertical lines either side of the mean. Asterisks (\**p* < .05) indicate the main effect of expectation on non-decision time (*t*_*0*_) and the interaction between emotion and expectation on drift rate (*v*; unexpected fearful > all other conditions).

We then applied model 7’s parameter optimisation approach to each participant’s response time data separately, to derive a *v* and *t*_*0*_ per participant, per condition. Model 7 provided adequate model fit across all participants (Kolmogorov-Smirnov test statistic: *p* = 0.696 ± 0.063, 0.575 to 0.851). We discovered that non-decision time (in seconds) was significantly shorter for expected (*M* = 0.882, 95% CI [0.782, 0.981]) than unexpected (*M* = 0.910, 95% CI [0.809 1.1011]) faces (*F*(1,30) = 9.352, *p* = .005, η_p_^2^ = 0.238), indicating that expectations hastened either stimulus encoding or response output processes (Ratcliff and McKoon, 2008). There was no significant main effect of emotion (*F*(1,30) = 0.036, *p* = .852, η_p_^2^ = .001) or interaction (*F*(1,30) = 0.801, *p* = .378, η_p_^2^ = .026).

For the *v* parameter, we applied a natural log transformation to correct the skewness of the data (original skewness: EN = 1.01, UN = 1.34, EF = 0.57, UF = 2.01, transformed skewness: EN = 0.41, UN = 0.56, EF = 0.14, UF = 0.94). We discovered that drift rate (in log units per second, towards the estimated threshold from relative starting point 0.5) was greater overall for fearful (*M* = 1.477, *95% CI* [1.366, 1.587]) than neutral (*M* = 1.394, *95% CI* [1.290, 1.498]) faces (*F*(1,30) = 8.122, *p* = .008, η _p_^2^ = 0.213), and was also greater overall for unexpected (*M* = 1.469, 95% CI [1.360, 1.578]) than expected (*M* = 1.402, *95% CI* [1.301, 1.503]) faces (*F*(1,30) = 13.200, *p* =.001, η _p_^2^ = 0.306). Critically, however, there was an interaction (*F*(1,30) = 5.933, *p* = .021, η _p_^2^ = 0.165), such that drift rate was only significantly increased for unexpected than expected *fearful* (difference: *M* = 0.122, *95% CI* [0.048, 0.197]) faces (*t*(30) = 3.351, *p* = 0.002, *p*_*bonf*_ = .004), while there was no significant difference between unexpected and expected *neutral* (difference: *M* = 0.013, *95% CI* [-0.027,0.052]) faces (*t*(30) = 0.648, *p* = .522, *p*_*bonf*_ = 1.00). These results indicate that evidence accumulated at a faster rate for unexpected fearful faces, relative to all other conditions.

Overall, the DDM results illustrate that the differential effect of prior expectations on response times to neutral and fearful faces may be explained by a combination of non-decision time and drift rate in the decision-making process. Non-decision time, representing stimulus encoding and/or motor response time, is hastened by prior expectations, explaining the faster response times to expected than unexpected neutral faces. For fearful faces, on the other hand, evidence accumulation is accelerated specifically for *unexpected* fearful faces; hence, the faster response times to unexpected than expected fearful faces.

### EEG reveals occipital, temporal, and frontal networks associated with emotional expectations

The DDM analysis explained the pattern of the response time via modulation of non-decision time and drift rate, such that response times to expected faces are accelerated (faster sensory encoding and/or motor execution) but response times to *fearful* faces are even faster when they are unexpected (more rapid evidence accumulation). We investigated this further by examining the timing of neural correlates with emotion and expectation processing. We conducted a General Linear Model in SPM to determine when and where in the scalp activity there was 1) a main effect of emotion (neutral vs. fearful), 2) a main effect of expectation, and 3) an interaction between emotion and expectation. This was achieved by conducting a full factorial ANOVA, with gender and block order added as covariates of no interest. After restricting the time window to 0 to 2 seconds post-stimulus onset (since we were interested in the activity *preceding* response times) and correcting for multiple comparisons (*p* < .05, cluster-level FWE), we did not observe significant differences in amplitude between expected and unexpected faces across the scalp. This was the case for both neutral and fearful faces, as there was no significant interaction effect. There were, however, three significant clusters for fearful versus neutral face activation (not shown). Both clusters spanned left occipital-parietal electrodes, where fearful faces elicited greater negative activity from 1.353 to 1.439 seconds and then from 1.661 to 1.768 seconds.

In conjunction with our scalp analysis, we also estimated the neural sources underlying the scalp activity from 0 to 2 seconds post-stimulus onset. We first looked at emotion and expectation effects and found significant main effects of each, as well as an interaction (see **Fig. 4**). There was significantly (*p* < .05, clusters FWE-corrected) greater activity for neutral than fearful faces in left V3/V4 and for fearful than neutral faces in the right middle temporal gyrus, in line with previous fMRI research (Sabatinelli et al., 2011). For expected faces, there was significantly greater activity in the left and right inferior frontal gyrus (IFG), left V3/V4, right middle frontal gyrus, and right temporal pole, supporting previous fMRI studies on expectation across sensory modalities (Hedge et al., 2015, Osnes et al., 2012). In contrast, there was significantly greater activity for unexpected faces in left V1/V2, consistent with previous fMRI work (Summerfield and Koechlin, 2008, Kok et al., 2012). Finally, the emotion by expectation interaction consisted of a greater surprise effect (that is, greater activity for unexpected than expected) for fearful than neutral faces in the right IFG and the left superior temporal gyrus (STG). In contrast, there was a greater surprise effect for neutral faces in the right middle temporal gyrus (MTG) and STG. These findings are consistent with previous fMRI studies demonstrating a role for the IFG in anticipating negative stimuli (Ueda et al., 2003, Sharot et al., 2011). Overall, our scalp and source analyses revealed a network of occipital, temporal, and frontal areas associated with emotion and expectation.

**Figure 4.**
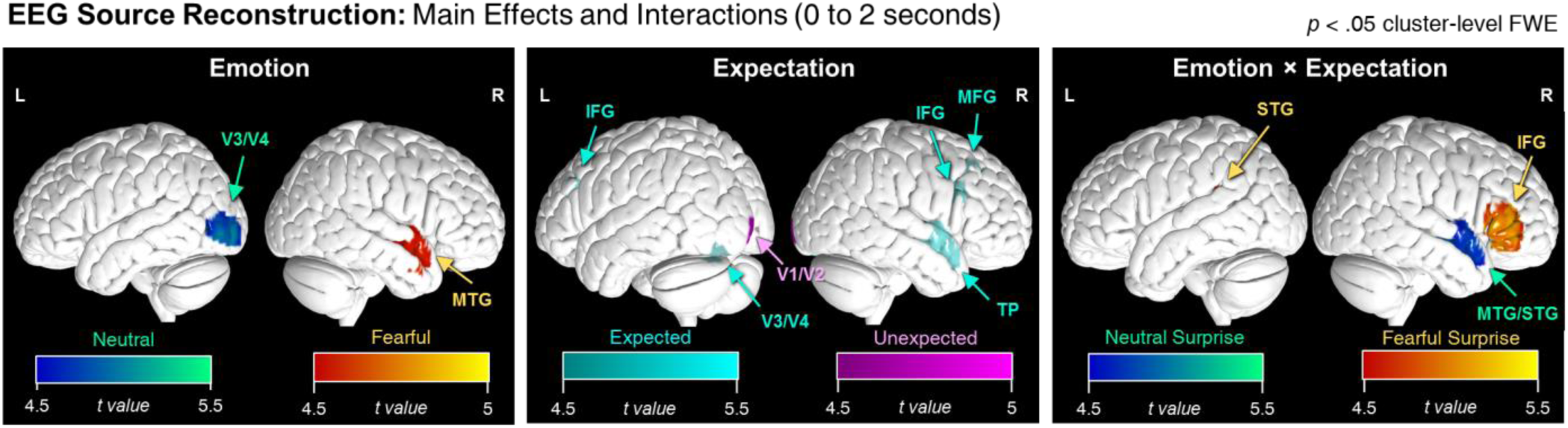
Source reconstruction reveals networks underlying emotion and expectation processing. The significant sources for emotion (left), expectation (middle), and the emotion by expectation interaction (right) are shown. Here, we show the *t* statistics for Neutral (minus Fearful), Fearful (minus Neutral), Expected (minus Unexpected), Unexpected (minus Expected), Neutral Surprise ([UN – EN] minus [UF – EF]), and Fearful Surprise ([UF – EF] minus [UN – EN]). L = left, R = right, MTG = middle temporal gyrus, IFG = inferior frontal gyrus, MFG = middle frontal gyrus, TP = temporal pole, IPL = inferior parietal lobule, STG = superior temporal gyrus. All clusters shown are *p* <.05 cluster-level FWE corrected.

### Higher drift rate for fearful surprise covaries with greater activity in the right IFG and faster non-decision time covaries with greater activity in visual areas

We then turned towards a model-based approach to EEG analysis to further elucidate how decision-making mechanisms relate to neural activity. This modeling analysis on response time revealed that drift rate was accelerated for fearful surprise and also that there was faster non-decision time for expected than unexpected faces. To find the neural correlates of these effects, we first conducted a scalp-level GLM similar to the one described above except that we also included non-decision time as a covariate of interest. We specifically investigated when and where non-decision time covaried with surprise-related neural activity (i.e. expected vs. unexpected). This revealed a cluster of central occipital activity from 1.106 to 1.188 seconds(not shown).

At the source level, we examined activity across three time windows (0.5 to 1 second, 1 to 1.5 seconds, and 1.5 to 2 seconds) to see whether there might be dynamic changes in significant sources over time. During only the 0.5 to 1 second window, we found that there was significantly greater activity for surprise in left V1/V2 for people who had faster non-decision time (*p* < .05, clusters FWE-corrected; see **Fig. 5**). Altogether, these results suggest that shorter non-decision time for expected than unexpected trials likely reflects faster stimulus encoding (Ratcliff and McKoon, 2008), supporting previous studies finding early spatiotemporal correlates (Nunez et al., 2017) and greater activity for surprise in the primary visual cortex (Summerfield and Koechlin, 2008, Kok et al., 2012).

**Figure 5.**
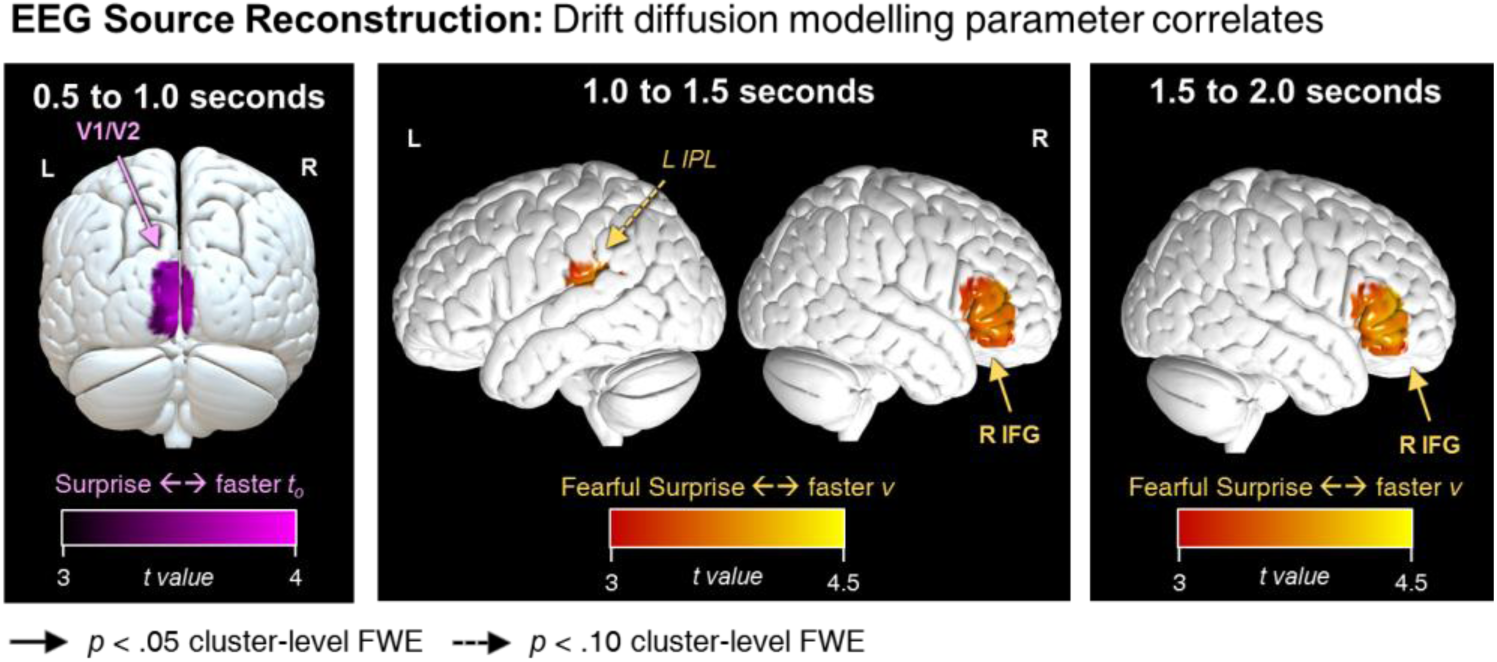
EEG source activity correlates with decision-making parameters. Estimated source activity is shown for time windows 0.5 to 1, 1 to 1.5, and 1.5 to 2 seconds post-stimulus onset. For each, neural correlates were investigated between faster non-decision time (*t*_*0*_) and neural activity for surprise (unexpected – expected) and between faster drift rate (*v*) and neural activity for fearful surprise ([unexpected – expected fearful faces] – [unexpected – expected neutral faces]). Clusters are thresholded at *p* < .10 FWE but note that the left V1/V2 and right IFG clusters are all significant at *p* < .05 FWE. Pink heat maps represent *t*-values for the surprise vs. *t*_*0*_ correlation and orange heat maps are for the fearful surprise vs. *v* correlation. L = left, R = right, V1/V2 = primary/secondary visual cortex, IFG = inferior frontal gyrus, IPL = inferior parietal lobe.

For the log-transformed drift rate parameter (*v*), we found that there was greater activity in the right IFG for fearful than neutral surprise when drift rate parameters were higher (see **Fig. 5**). This was the case for time windows spanning 1 to 2 seconds. There was also a borderline-significant cluster in the left inferior parietal lobule (*p* = .067) within the 1 to 1.5 second time window. Hence, the right IFG appears to be associated with accelerated evidence accumulation to surprising threats emerging into conscious perception. Note that we did not observe any significant clusters of activity at the scalp level after correcting for multiple comparisons (*p* < .05 cluster-level FWE) for the correlation between drift rate and the emotion × expectation interaction (that is, a greater effect of surprise for fearful than neutral faces).

Overall, this model-based neuroimaging analysis revealed the evolution of source activity over time for each of our key DDM parameters and how these covaried with emotion-induced and expectation-induced neural activity. Particularly, we found an initial, short-lived increase in primary visual cortex activity for surprise when non-decision time was faster. Finally, there was a consistently greater effect of fearful than neutral surprise when drift rate was higher in the right IFG from 1 to 2 seconds, suggesting a potential role for this area in facilitating earlier conscious perception of unexpected threats.

## DISCUSSION

Here we set out to explore the interaction between emotion and expectation on the conscious awareness of stimuli, and to determine the neural mechanisms underlying these effects. To achieve this, we modelled behavioural responses to neutral and fearful faces that emerged from continuous flash suppression (CFS) and were either expected or unexpected, and correlated the resulting parameters to human neural activity recorded with EEG. In line with previous research, we found that expectation accelerated the conscious perception of faces. Model-based EEG analyses revealed that this was driven by faster non-decision time for expected than unexpected faces, which correlated with increased activation of early visual cortex shortly after stimulus onset. Fearful faces were fastest to be consciously perceived overall but, crucially, were either unaffected by expectation (Experiment 1) or were faster to break through into consciousness when unexpected (Experiment 2). We discovered that this was driven by an especially fast rate of evidence accumulation (drift rate) for surprising fearful faces, which correlated with sustained activity in the right IFG. These results are consistent with the **Survival Hypothesis**, and suggest that occipital and frontal networks in the human brain facilitate the fast detection of danger, even when such threats are improbable and tangential to the task at hand.

Our results provide experimental evidence for the folk notion that ‘we see what we want to see’. Fearful faces, which present an evolutionarily-relevant ambiguous threat signal, were more quickly detected than neutral faces, in line with previous research (Hedger et al., 2014, Capitão et al., 2014, Yang et al., 2007, Tsuchiya et al., 2009). Our analysis also revealed that fearful faces evoked stronger activity in the right MTG and accelerated the rate of evidence accumulation (drift rate), the latter of which supports previous research showing that emotional content increases drift rate in perceptual decision-making (Tipples, 2015) even if unconsciously-presented via CFS (Lufityanto et al., 2016).

Our results also support the folk notion that ‘we see what we expect to see’. Consistent with previous literature (Pinto et al., 2015, Hesselmann et al., 2010, Vetter et al., 2014, Hohwy et al., 2008), expected stimuli were detected faster than unexpected stimuli. Drift-diffusion modelling of reaction time data revealed that this effect was underpinned by a reduction in non-decision time with expectation (for both neutral and fearful faces), a finding that is consistent with previous research on temporal expectations (Jepma et al., 2012). Subsequent model-based EEG analyses revealed that this effect correlated with greater activity in left V1/V2 for surprise in only the earliest analysed time window (0.5 to 1 second post-stimulus onset), complementing a previous DDM-based EEG study on attention that found early (150-275ms post-stimulus onset) activity related to non-decision time (Nunez et al., 2017). Together, the early timing and location of these effects suggest that expectation accelerates stimulus encoding rather than motor execution, the correlates of which would likely occur in motor areas and much closer to the participants’ response (average response time ∼1.8 seconds). Consistent with this interpretation, recent studies have suggested that expectation ‘sharpens’ sensory representations in the primary visual cortex (Kok et al., 2012), or, alternatively, neural scaling (Alink et al., 2018), either of which could account for the reduced neural activity in response to expected faces that we observed here.

We discovered that expectation and emotion interacted to influence conscious perception, such that expectation hastened the detection of neutral but not fearful faces. Drift-diffusion modelling of reaction time data from Experiment 2 revealed that this interaction was driven by a faster rate of evidence accumulation (drift rate) for unexpected fearful faces, relative to all other conditions. This interaction effect could explain why previous studies using only neutral stimuli have found no influence of expectation on drift rate (De Loof et al., 2016). Similar effects on drift rate have previously been seen with attention (Tavares et al., 2017, Nunez et al., 2017), and previous work on inattentional blindness has suggested that unexpected threatening stimuli (e.g. spiders, guns or snakes:

New and German, 2015, Wiemer et al., 2013, Gao and Jia, 2017) draw more attention and thus are more likely to be noticed than unexpected neutral stimuli (but see also (Calvillo and Hawkins, 2016, Beanland et al., 2017). In accordance with this literature, we speculate that the enhanced drift rate to unexpected fearful faces and their subsequent early conscious perception might reflect an underlying interaction between (exogenous) attention and prediction. Consistent with this interpretation, a recent discussion of predictive coding theory suggests that attention interacts with prediction to optimise the expected precision of predictions via gain-modulation of prediction errors (Feldman and Friston, 2010), and a recent study conducted by our group provided experimental evidence for this theory (Smout, Tang, Garrido & Mattingley, 2019). Since prediction errors (in predictive coding) and drift rate (in sequential sampling) can be considered to be equivalent (under certain simplifying assumptions, see Bitzer et al., 2014), it follows from this theory that attended and unexpected stimuli should exhibit increased drift rate (or, equivalently, prediction errors), as we observed here.

Our source analysis revealed that the right IFG showed persistently greater signal for fearful than neutral surprise when drift rate was higher. This is consistent with findings for the involvement of the IFG in generating the mismatch negativity ERP response to unexpected stimuli (Garrido et al., 2009, Doeller et al., 2003, Opitz et al., 2002, Kim, 2014). Critically, our results further extend this finding by demonstrating that this effect is enhanced for threatening stimuli and is associated with accelerated evidence accumulation even while stimuli are breaking through into conscious perception. Hence, the right IFG may play a significant role in increasing the gain of prediction error signals for fearful faces. Indeed, previous research has found the IFG to respond more to fearful than neutral faces (Ishai et al., 2004, Luo et al., 2007), with IFG activity being predictive of fearful face perception near the threshold of conscious awareness (Pessoa and Padmala, 2005). Our time-resolved source-level GLM results suggest that unexpected threat triggers rapid evidence accumulation for dichoptically-suppressed face stimuli, involving the right IFG as early as 1 second (see **Fig. 5**) before a conscious perceptual decision is made. Future research could broaden the extent of this network by using MEG or fMRI to tap into both subcortical and cortical sources to see whether circuits including the amygdala (Tamietto and De Gelder, 2010, Mitchell and Greening, 2012) may contribute to unexpected threat responses in the IFG.

Overall, our results present a newly-discovered interaction between prior expectations and emotional expression that modulates how early we can make conscious perceptual decisions about faces. This effect was driven by an acceleration of early stimulus encoding by prior expectations, as well as an early and sustained increase in evidence accumulation, involving the right IFG, specifically for unexpected fearful faces. Although we took a measure of non-clinical state and trait anxiety that did not significantly correlate with behavioural or neural responses in our healthy participants, it is conceivable that the conscious perception of threat might be modulated by prior expectations in clinical anxiety, which is characterised by threat overexpectancy (Aue and Okon-Singer, 2015). It has also recently been shown that people with schizophrenia have aberrant expectations for threat (Dzafic et al., 2018, Barbalat et al., 2012). Hence, future computational psychiatric research may yield invaluable findings by exploring this line of research in people with various types of clinical anxiety (e.g. social anxiety, specific phobia, or post-traumatic stress disorder) or schizophrenia.

