## Supplementary Materials for "Surprising Threats Accelerate Evidence Accumulation for Conscious Perception"

### Experimental Setup

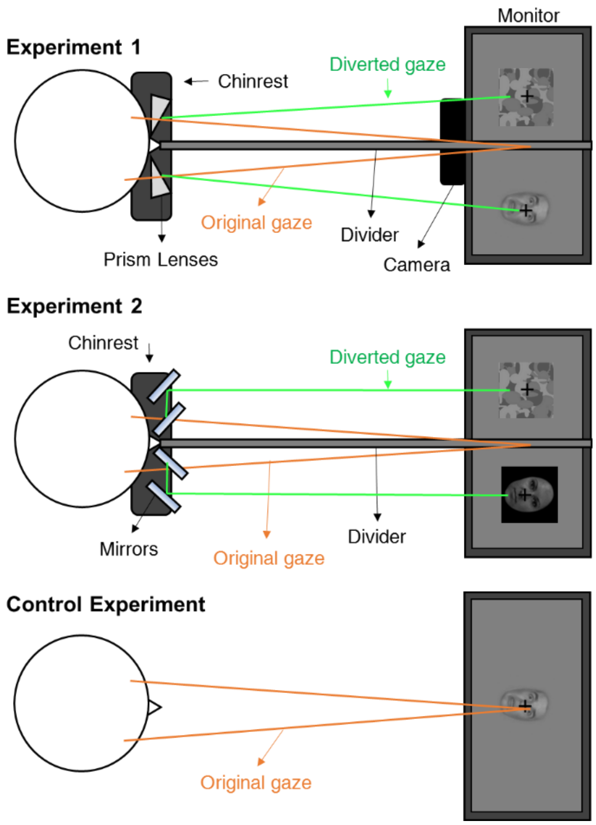


**Supplementary Figure 1. Equipment setup.** A bird's eye view is shown for the set-up for each of the three experiments. In Experiment 1, the participant viewed dichoptically-presented stimuli (Mondrian mask in dominant eye, face in non-dominant eye) through prism lenses. The set-up for Experiment 2 was the same as Experiment 1 but using stereoscopic mirrors instead of prism lenses and without a camera. The background of the face stimuli was also made black so as to enhance visually-evoked EEG responses. The Control Experiment used the same stimuli as Experiment 1 but participants simply viewed face stimuli without a mask or dichoptic presentation.

### Control Experiment

In a separate control experiment, we removed the mask from the bCFS paradigm to investigate whether the effects observed in Experiment 1 could occur without ambiguity introduced by perceptual suppression. Hence, the design of the control experiment was identical to Experiment 1 except that the face was presented to both eyes, without the mask.

#### *Participants*

We recruited 30 participants through the University of Queensland's Participation Scheme. Our sample consisted of 8 males and 22 females aged between 18 and 33 years ($M$ = 22.87, $SD$ = 2.47). All participants had normal or corrected-to-normal vision. Participants were compensated $20 AUD per hour for their time and provided written consent. This study was approved by the University of Queensland's Medical Research Ethics Committee.

#### *Stimuli and procedure*

The method was identical to Experiment 1 except that face stimuli were presented centrally and no prism lenses or mirrors were used for dichoptic presentation. Hence, we did not use an eye tracker to monitor whether both eyes were open and no headrest was used (see **Supplementary Fig. 1**). Also, we expected response times to be faster in this unsuppressed version of the task and so we increased the ITI by 1 s, giving a jittered ITI of 1.5 s to 2 s in steps of 100 ms, so that the time between motor responses would be more comparable with Experiments 1 and 2.

#### *Behavioural analysis*

We removed slow responses more than five standard deviations above the mean. Average trial counts (correct responses only) for expected faces were approximately 291 for neutral (275 to 298) and fearful (272 to 299) expressions and, for unexpected faces, 58 (54 to 60) for neutral and 59 (53 to 60) for fearful. We entered the trial data from each participant into a LME analysis akin to those described in Experiments 1 and 2.

#### *Results*

The mean response times were 0.898 s ($SD$ = 0.438 s) for expected neutral, 0.905 s ($SD$ = 0.418 s) for unexpected neutral, 0.903 s ($SD$ = 0.437 s) for expected fearful, and 0.913 s ($SD$ = 0.484 s) for unexpected fearful faces. The results of the likelihood ratio tests revealed that none of our models were significantly more likely than the null model (log likelihoods, minus the null model: emotion = -1.1, expectation = -1.0, emotion + expectation = 0.4, interaction = -0.4), indicating that there were no meaningful differences between our experimental conditions. Therefore, we can conclude that the interocular suppression between the mask and the faces was necessary for emotion and expectation to influence response times in our experimental paradigm.
